# Choanoflagellates and the ancestry of neurosecretory vesicles

**DOI:** 10.1101/2020.05.24.111997

**Authors:** Ronja Göhde, Benjamin Naumann, Davis Laundon, Cordelia Imig, Kent McDonald, Benjamin H. Cooper, Frederique Varoqueaux, Dirk Fasshauer, Pawel Burkhardt

## Abstract

Neurosecretory vesicles are highly specialized trafficking organelles important for metazoan cell-cell signalling. Despite the high anatomical and functional diversity of neurons in metazoans, the protein composition of neurosecretory vesicles in bilaterians appears to be similar. This similarity points towards a common evolutionary origin. Moreover, many key neurosecretory vesicle proteins predate the origin of the first neurons and some even the origin of the first animals (metazoans). However, little is known about the molecular toolkit of these vesicles in non-bilaterian metazoans and their closest unicellular relatives, making inferences about the evolutionary origin of neurosecretory vesicles extremely difficult. By comparing 28 proteins of the core neurosecretory vesicle proteome in 13 different species, we demonstrate that most of the proteins are already present in unicellular organisms. Surprisingly, we find that the vesicle residing SNARE protein synaptobrevin is localized to the vesicle-rich apical and basal pole in the choanoflagellate *Salpingoeca rosetta*. Our 3D vesicle reconstructions reveal that the choanoflagellates *Salpingoeca rosetta* and *Monosiga brevicollis* exhibit a polarized and diverse vesicular landscape. This study sheds light on the ancestral molecular machinery of neurosecretory vesicles and provides a framework to understand the origin and evolution of secretory cells, synapses, and neurons.

## Introduction

Coordinated cell-cell signalling is one of the most significant pre-requisites for the evolution of multicellular organisms. In metazoans, a variety of highly specialized neuronal cell types has evolved facilitating signal perception, integration, propagation, offset and response building up complex neuronal circuits (Schmidt-Rhaesa *et al*. 2015). In these neuronal circuits, signals are often propagated in a directional manner via several cells at distinct cell-cell contact sites called synapses. While metazoan synapses have been extensively studied through several decades their evolutionary origin is still unresolved (Ryan and Grant 2009, Nickel 2010, Burkhardt and Sprecher 2017, Varoqueaux and Fasshauer 2017, Arendt 2020).

To elucidate the evolution of synapses in metazoans, it is necessary to investigate the presence and ancestral function of key synaptic components in related unicellular organisms. One of these key components are neurosecretory vesicles that deliver neuroreactive cargos between pre- and postsynaptic cells. At least two different types of neurosecretory vesicles, synaptic vesicles (SVs) and dense-core vesicles (DCVs) are involved in regulated secretion in neurons, but also in endocrine and neuroendocrine cells (Morgan and Burgoyne 1997, Hannah *et al*. 1999). Synaptic vesicles concentrated at pre-synapses facilitate signal propagation via the fusion of the vesicular membrane with the pre-synaptic plasma membrane and neurotransmitter release into the synaptic cleft. In endocrine and neuroendocrine cells, but also in neurons, dense-core vesicles function in multiple biological processes via the release of proteins or neuropeptides. In neurons, dense-core vesicles are found in many different parts of the cell, including dendrites, varicosities and synaptic terminals, where they fulfil important roles for synaptic transmission, memory formation and neuronal survival (Scalettar 2006). The protein compositions of these two classes of neurosecretory vesicles are well characterized (Takamori *et al*. 2006, Wegrzyn *et al*. 2007, Wegrzyn *et al*. 2010). Despite different cargos and biological roles, the protein composition of neurosecretory vesicles appears to be similar (Grønborg *et al*. 2010) and are composed of a set of core proteins, which can be assigned to specific categories (Figure 1) (adapted from (Jahn and Boyken 2013)).

**Figure. 1.**
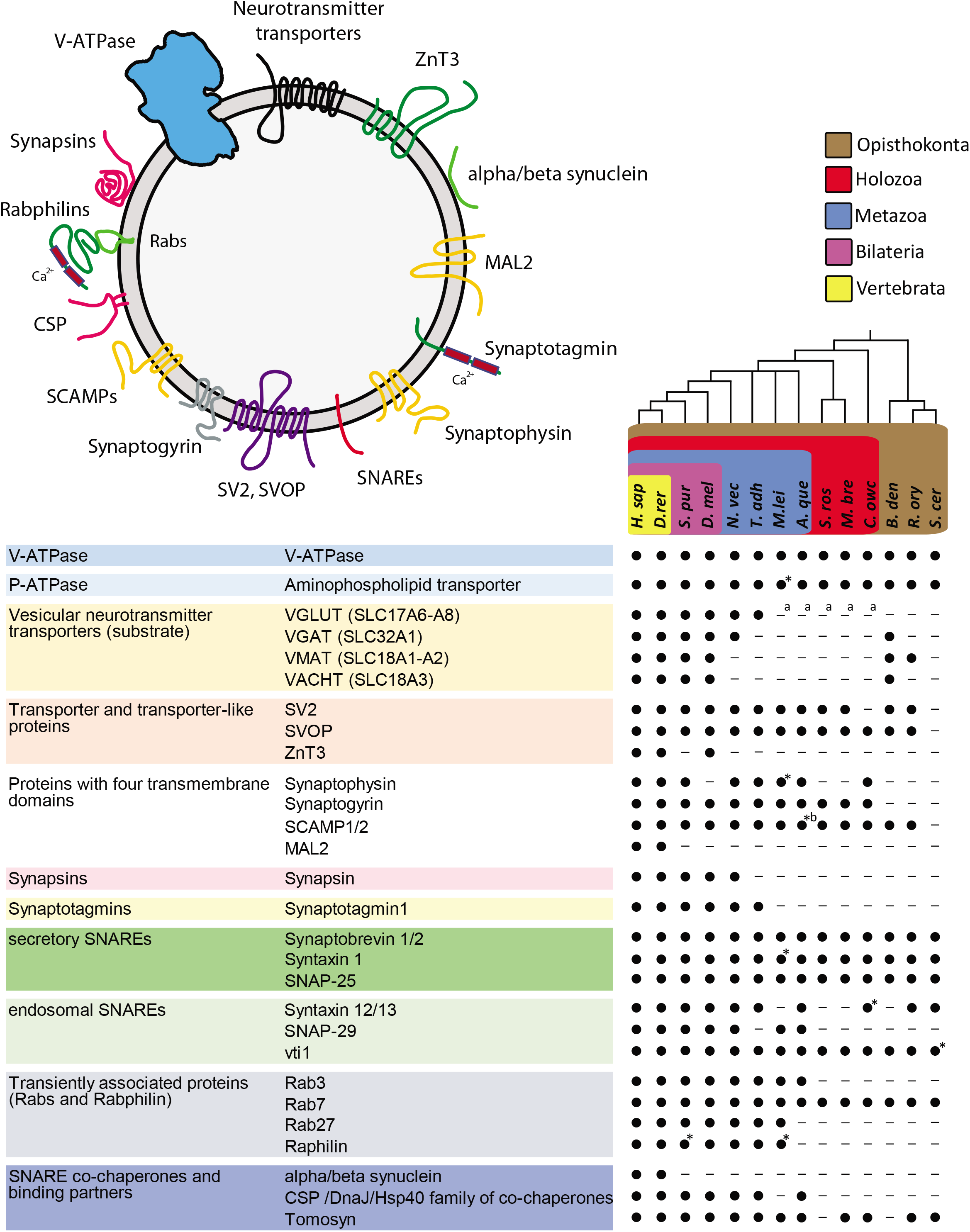
Neurosecretory vesicle proteins in metazoans and their closest living relatives. (Top) Schematic model of the core molecular components of metazoan neurosecretory vesicles. (Below) Core proteins of metazoan neurosecretory vesicles can be assigned to eleven categories: V-ATPases, vesicular neurotransmitter transporters, transporter and transporter-like proteins, proteins with four transmembrane domains, synapsins, synaptotagmins, secretory SNAREs, endosomal SNAREs, transiently associated proteins SNARE binding partners and co-chaperones (modified after Jahn and Boyken 2013). Black dots indicate the presence of clear protein sequence homologs (also see supplementary table 1), while lines indicate that a homolog was not detected in the respective organism. Taxonomic groupings are indicated as follows: Brown box, Opisthokonta; red box, Holozoa; blue box, Metazoa; violet box, Bilateria; yellow box, Vertebrata. Phylogenetic tree based on a consensus phylogeny (Baldauf 2003, Ruiz-Trillo *et al*. 2008, Philippe *et al*. 2009, Fairclough *et al*. 2013). A. que, *Amphimedon queenslandica*; B. den, *Batrachochytrium dendrobatidis*; C. ele, *Caenorhabditis elegans*; C. owc, *Capsaspora owczarzaki*; D. mel, *Drosophila melanogaster*; H. sap, *Homo sapiens*; M. bre, *Monosiga brevicollis*; M. mus, *Mus musculus*; N. vec*, Nematostella vectensis*; O. car, *Oscarella carmela*; R. ory, *Rhizopus oryzae*; S. cer, *Saccharomyces cerevisae*; S. pur, *Strongylocentrotus purpuratus*; S. ros, *Salpingoeca rosetta*; T. adh, *Trichoplax adhaerens*. * = protein of interest-like, ^a^ = putative SLC17A5-homolog, ^b^ = domain structure lost.

Choanoflagellates are the closest unicellular relatives of metazoans and exhibit a surprisingly rich repertoire of neuronal protein homologs (King *et al*. 2008, Alié and Manuel 2010, Fairclough *et al*. 2013, Burkhardt *et al*. 2014, Yang *et al*. 2015, Hoffmeyer and Burkhardt 2016). Particularly interesting is the recent observation of morphologically distinct intracellular vesicle populations (Laundon *et al*. 2019) and the presence of plasma membrane contacts between colonial cells in the choanoflagellate *S. rosetta* (Naumann and Burkhardt 2019). These observations make choanoflagellates a suitable model to investigate the ancestry of neurosecretory vesicles and the evolutionary origin of metazoan synapses. In the present study, we performed a comparative analysis of neurosecretory vesicle proteins together with a morphological characterization of the vesicle types in *S. rosetta* and *M. brevicollis*. Using comparative cross-species protein analysis in combination with immunohistochemistry, serial ultrathin transmission electron microscopy (ssTEM) and 3D-reconstruction, we show that choanoflagellates exhibit a rich repertoire of neurosecretory vesicle proteins and a polarized and diverse vesicular landscape.

## Results

### a) Comparative analysis reveals the ancestry of neurosecretory vesicle proteins

Neurosecretory vesicles are composed of a “core proteome” which can be subdivided into specific categories: ATPases, transporters and transporter-like proteins, proteins with four transmembrane domains, synapsins, synaptotagmins, SNAREs, SNARE co-chaperones and SNARE binding partners, as well as proteins that are only transiently associated with synaptic vesicles (Figure 1). Based on this core proteome, we selected 28 proteins with at least on representative from each category to perform a survey for respective homologs. This survey was conducted in a total of 13 different eukaryotic species (*Danio rerio, Strongylocentrotus purpuratus, Drosophila melanogaster, Nematostella vectensis, Trichoplax adhaerens, Mnemiopsis leidyi, Amphimedon queenslandica, Salpingoeca rosetta, Monosiga brevicollis*, *Capsaspora owczarzaki*, *Batrachochytrium dendrobatidis*, *Rhizopus oryzae*, *Saccharomyces cerevisiae* (Figure 1)).

Overall, we found that ~39% of the examined neurosecretory vesicle proteins are restricted to metazoans. For example, synapsin, one of the most abundant synaptic vesicle proteins (Takamori *et al*. 2006), the synaptic-associated zinc transporter ZnT3 (Palmiter *et al*. 1996), the calcium sensor synaptotagmin1 (Yoshihara *et al*. 2003), the co-chaperone cysteine string protein (CSP) (Tobaben *et al*. 2001), MAL2 (Grønborg *et al*. 2010) and synuclein (Uéda *et al*. 1993) are only found in metazoans. Strikingly, and in accordance with previous studies (Gogarten *et al*. 1989, Gogarten *et al*. 1992, Axelsen and Palmgren 1998, Mackiewicz and Wyroba 2009, Burkhardt *et al*. 2014, Moroz and Kohn 2015), our results show, that the majority (~61%) of the examined neurosecretory vesicle proteins are also present in unicellular eukaryotes (Figure 1), as we found secretory SNAREs, Rab7, V-and P-ATPase protein sequences in all investigated organisms. We also identified the “four transmembrane domain protein” synaptophysin in the unicellular eukaryote *C. owczarzaki*, as well as synaptogyrin in *C. owczarzaki*, and in the two choanoflagellate species *M. brevicollis* and *S. rosetta*. In addition, we found the synaptic vesicle protein 2 (SV2) in most of the investigated organisms and the SV2-related protein (SVOP) in all species, except for *S. cerevisiae*. The analysed neurotransmitter transporters showed diverse presence and absence patterns for the different species. While we found vesicular glutamate transporters (VGLUT) in all investigated bilaterians, *N. vectensis* and *T. adhaerens*, we only found members of vesicular monoamine transporters (VMAT) and vesicular acetylcholine transporters (VAChT) in bilaterians and in some of the investigated fungi. The vesicular GABA transporter (VGAT) was found in all investigated bilaterians, *N. vectensis* and in the fungus *B. dendrobatidis*. However, this transporter seems to be absent in *T. adhaerens*, *M. leidyi*, *A. queenslandica, S. rosetta*, *M. brevicollis*, *C. owczarzaki, R. oryzae* and *S. cerevisiae*.

Our comparative analysis revealed that ~61% of the core neurosecretory vesicle proteins evolved prior to the emergence of the first metazoans. To further assess the evolutionary origin of neurosecretory vesicles, we analysed the localization of the vesicle-associated protein synaptobrevin in the choanoflagellate *S. rosetta*.

### b) Synaptobrevin as a putative secretory vesicle marker in the choanoflagellate *Salpingoeca rosetta*

The vesicle-associated SNARE protein synaptobrevin 1/2 (VAMP 1/2), together with Syntaxin 1 and SNAP-25 forms a stable complex which mediates the fusion of neurosecretory vesicles with the pre-synaptic plasma membrane and thereby release neurotransmitters/peptides in a variety of different metazoans (Kloepper *et al*. 2008). The genome of the choanoflagellate *S. rosetta* encodes for a single synaptobrevin. *S. rosetta* synaptobrevin contains a highly conserved coiled-coil region responsible for SNARE complex formation (Lin and Scheller 1997, Sutton *et al*. 1998) (Figure 2A and B) and a single C-terminal transmembrane domain (Figure 2A). *S. rosetta* synaptobrevin displays sequence identity to human synaptobrevin 1 of 38% and to human synaptobrevin 2 of 36%.

**Figure 2.**
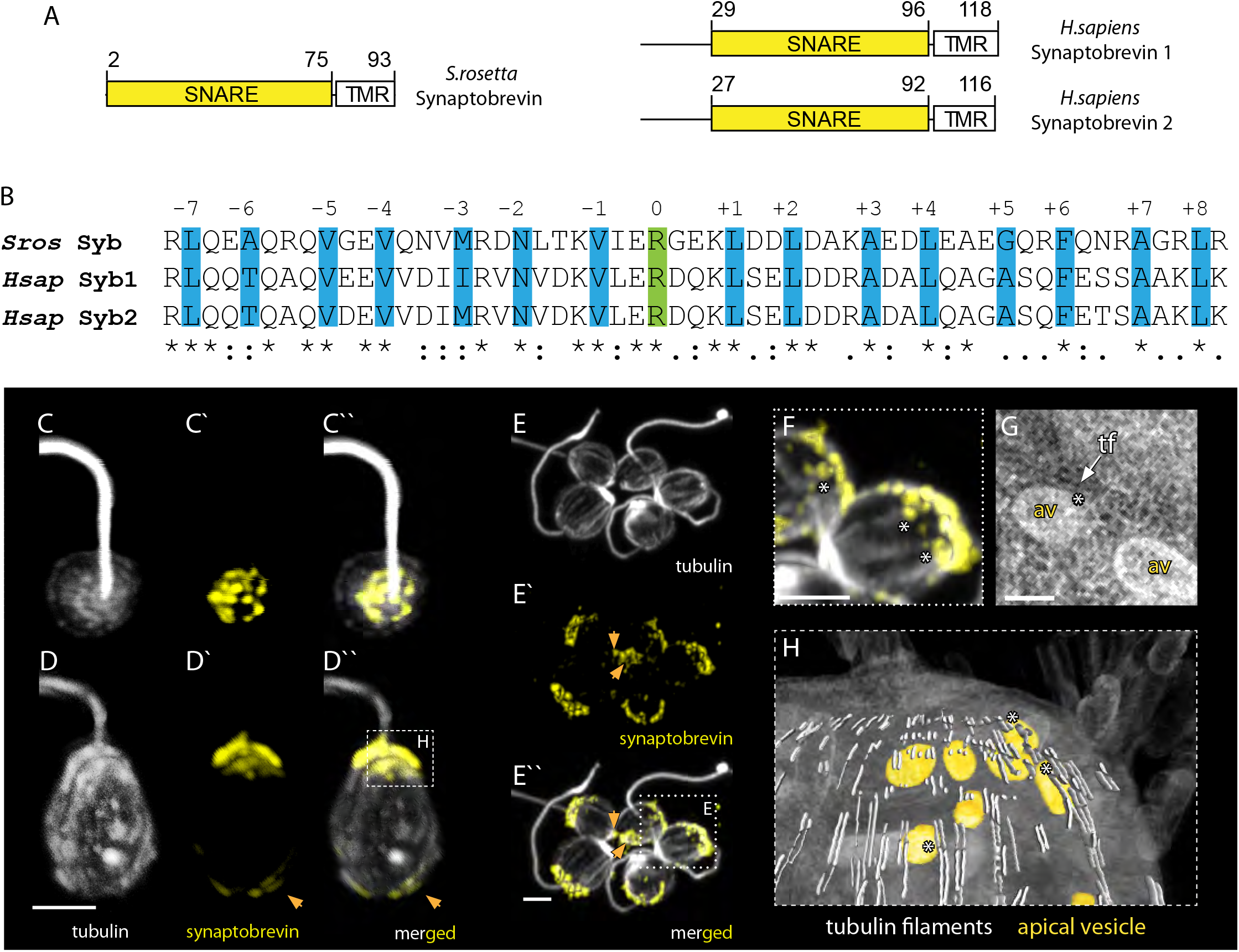
Synaptobrevin in the choanoflagellate *Salpingoeca rosetta*. A, Domain architecture of *Salpingoeca rosetta* synaptobrevin and *Homo sapiens* synaptobrevin 1 and 2. B, Sequence alignment of the SNARE motif of *S. rosetta* synaptobrevin and *H. sapiens* synaptobrevin 1 and 2. The 16 layers (highlighted in blue including layers −1 to −7 and layers +1 to +8) important for SNARE complex formation are shown. The conserved arginine residues forming the ionic 0 layer is shown in green. CC’’, apical view of an *S. roesetta* cell stained with antibodies against tubulin (grey) and synaptobrevin (yellow). C, tubulin. C’, synaptobrevin. C’’, merged. D-D’’, lateral view of a different *S. rosetta* cell stained with antibodies against tubulin and synaptobrevin. D, tubulin. D’, synaptobrevin. D’’, merged. The dashed square in D’’ indicates to position of H. E-E’’, a rosette colony of *S. rosetta* stained with the same antibodies as in C. The orange arrows indicate to a basal synaptobrevin signal. E, tubulin. E’, synaptobrevin. E’’, merged. The dotted square in E’’ indicates to position of F. F, Synaptobrevin-positive vesicles are in close contact with tubulin-positive cytoskeletal filaments. F, TEM image showing the close contact between apical vesicles and tubulin filaments. G, 3D reconstruction of the apical region of an *S. rosetta* cell. Apical vesicles (av) are colored in orange, tubulin filaments (tf) in light grey and the soma in half-transparent grey. Close contact of vesicles and cytoskeletal filaments are indicated with a white asterisk. The scale bar is 1 μm.*Sros, Salpingoeca rosetta; Hsap, Homo sapiens*.

To assess the subcellular localization of synaptobrevin in *S. rosetta* by immunostaining, we raised a polyclonal antibody against the soluble portion of the protein (aa 1-75). Our immunohistochemical staining experiments revealed synaptobrevin localization predominantly to the apical part of the cell (Figure 2C-F), confirming previous results we obtained in *M. brevicollis* (Burkhardt *et al*. 2011). Surprisingly, we also noticed synaptobrevin localization at the basal part of the cell, suggesting the presence of secretory vesicles on two opposing sites of the cell. A very similar staining pattern was detected in *S. rosetta* cells of rosette colonies (Fig. 2E). Additionally, an overlap of the tubulin signal with single synaptobrevin positive vesicles can be detected (Figure 2E-F). This co-association of cytoskeletal (tubulin) filaments and at least apical vesicles is also present in TEM sections revealed by 3D reconstruction (Figure 2G and H). We did not detect synaptobrevin signals at putative plasma membrane contact sites, predominantly located in the median area of cell somata (Fig. 2E’’).

### c) Diverse and polarized vesicular landscape in choanoflagellates

To investigate the number and diversity of vesicles in choanoflagellates we reconstructed the vesicular landscape in unicellular *M. brevicollis* (Figure 3A, supplementary video 1). We were able to identify 163 vesicles in total which we assigned to five different gross vesicle types. (1) Electron-dense Golgi-associated vesicles (N = 79; mean diameter 54 nm) are in the apical region of the cell, close to the Golgi apparatus (Figure 3B, B’). In the reconstructed cell, a tubulus of the endoplasmic reticulum (ER) is located basally to the Golgi apparatus. Vesicles between this ER tubulus and Golgi cisternae exhibit the same size as apical Golgi-associated vesicles but are more heterogeneous regarding their electron density (Figure 3B’). However, they often appear slightly more electron-lucent compared to apical golgi-associated vesicles. (2) Small electron-lucent vesicles, resembling the vesicles between the Golgi-apparatus and ER tubulus but slightly larger (N = 51; mean diameter 72 nm), can be found in the whole cell soma with a higher concentration in the basal area (Figure 3C, C’). (3) Apical vesicles (N = 6; mean diameter 116 nm) are present in low numbers. They exhibit a higher electron density and are located in close proximity to the apical complex (Figure 3 D, D’). (4) Large extremely electron-lucent vesicles (N = 15; mean diameter 129 nm) are present in a scattered pattern within the whole cell soma (Figure 3E, E’). (5) Large electron-dense vesicles (N = 12; mean diameter 129 nm) are present mainly in the basal third of the cell soma (Figure 3 F, F’).

**Figure 3.**
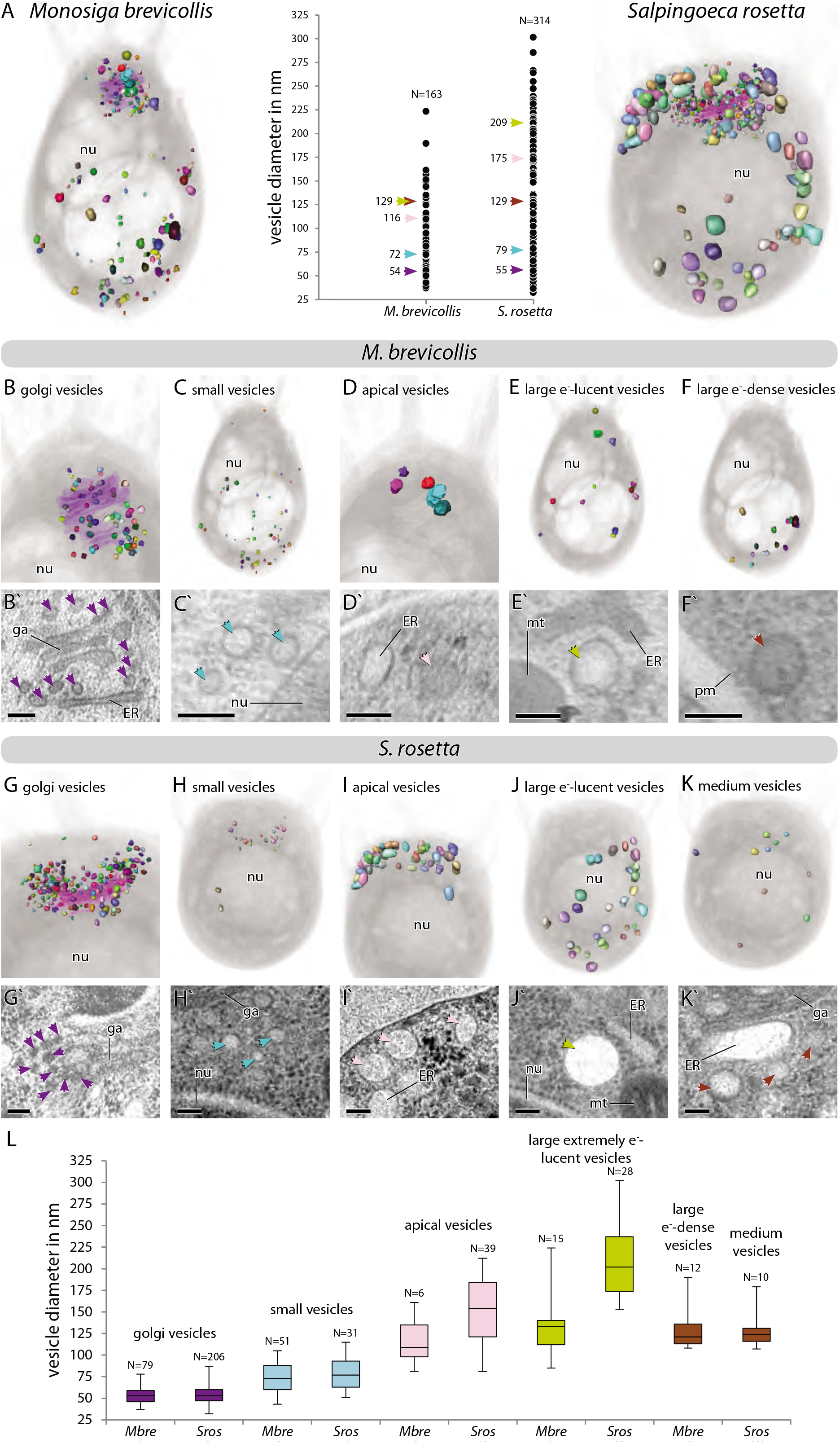
The diverse vesicular landscape of choanoflagellates. A, 3D reconstruction of all vesicles in *M. brevicollis* (right) and *S. rosetta* (left). Individual vesicles are coloured randomly, and the cell is shown in half-transparent grey. A plot of all vesicle diameters measured is given in the middle. Mean diameters of different vesicle types are indicated by arrows in the same colours as in B-L. B-F, Visualization of separated vesicles of each vesicle type to show the localization within the soma of *M. brevicollis*. The Golgi-apparatus is shown in half-transparent lilac in B. TEM image showing each vesicle type is given beneath the 3D model (B’-F’). G-K, Visualization of separated vesicles of each vesicle type to show the localization within the soma of *S. rosetta*. The Golgi-apparatus is shown in half-transparent lilac in B. TEM image showing each vesicle type is given beneath the 3D model (G’-K’). L, Box and whiskers plot of the vesicle diameters within the different vesicle types (also see supplementary table 2 and supplementary video 1 and 2). *Mbre, Monisiga brevicollis; Sros, Salpingoeca rosetta*. nu = nucleus, ga = Golgi apparatus, ER = endoplasmic reticulum, mt = mitochondria, pm = plasma membrane.

To compare our findings with another choanoflagellate species we reconstructed the vesicular landscape of unicellular *S. rosetta* (Figure 3A, supplementary video 2). We were able to identify 314 vesicles in total which we assigned to five different gross vesicle types. (1) Electron-dense Golgi-associated vesicles (N = 206; mean diameter 55 nm) are located in the apical region of the cell close to the Golgi apparatus (Figure 3G, G’). (2) Small electron-lucent vesicles (N = 31; mean diameter 79 nm) can be found mainly in the apical region (Figure 3H, H’). They resemble the Golgi-associated vesicles located between the basal Golgi apparatus and the ER tubules in *M. brevicollis* (Figure 3B, B’). However, the small vesicles in *S. rosetta* are very homogenous regarding their electron density and therefore also resemble the small vesicles of *M. brevicollis* (Figure 3C, C’). (3) Larger apical vesicles (N = 39; mean diameter 175 nm) are located in close proximity to the apical complex (Figure 3I, I’). These vesicles are more electron-lucent and often more ovoid compared to the apical vesicles in *M. brevicollis*. (4) Large extremely electron-lucent vesicles (N = 28; mean diameter 209 nm) are present within the whole cell soma (Figure 3J, J’). (5) A few medium-sized, electron-lucent vesicles (N = 10; mean diameter 129 nm) are present in a scattered pattern (Figure 3 F, F’).

### d) Diversity of vesicles in two choanoflagellate species – commonalities and differences

A comparison between the vesicular landscapes of *M. brevicollis* and *S. rosetta* reveals differences in vesicle numbers of similar types as well as of different vesicle types (Figure 3L). Golgi-associated vesicles are of similar mean diameter in *S. rosetta* and *M. brevicollis*. However, *S. rosetta* exhibits 2.6 times more Golgi-associated vesicles than *M. brevicollis*. The mean diameter of small vesicles is also similar but slightly larger in *S. rosetta*. These vesicles differ in their abundance (1.6 times more in *M. brevicollis*) and cellular localization (compare Figure 3C and H). Apical vesicles are different regarding their number (6.5 times more in *S. rosetta*), mean diameter (1.5 times larger in *S. rosetta*) and form (spherical in *M. brevicollis* and spherical to ovoid in *S. rosetta*). Similar differences can be observed for large (extremely) electron-lucent vesicles regarding their number (1.9 times more in *S. rosetta*) and mean diameter (1.6 times larger in *S. rosetta*). Large electron-dense vesicles of *M. brevicollis* show no similarities compared to the medium vesicles in *S. rosetta* and might represent different vesicle types.

## Discussion

In agreement with previous studies (Ryan *et al*. 2009, Burkhardt *et al*. 2014, Moroz *et al*. 2015, Liebeskind *et al*. 2017, Abrams and Sossin 2019), our results show that many components of the core proteome of neurosecretory vesicles have a pre-metazoan origin. In addition, we discovered, that some of the vesicular transporters may be even older than previously thought due to their presence in the fungus *B. dendrobatidis*. Furthermore, we showed the presence of a diverse vesicular landscape in choanoflagellates, the closest unicellular relatives of metazoans. Some of the vesicle types are closely associated with cytoskeletal components (tubulin filaments) and seem to be concentrated either apically or basally, indicating the presence of a directed vesicular transport system in choanoflagellates. Neurosecretory vesicles, concentrated either apically or basally and closely associated with tubulin filaments are observed in a variety of metazoan neuronal cell types as well (Squire *et al*. 2012). The observed similarities in vesicular protein composition and landscape in choanoflagellates and metazoan nerve cells could be explained by at least two different hypotheses:

1. Secretion-based cell-cell signalling, employing components of metazoan synaptic transmission, was already present in the last common ancestor of the holozoans. There, it could have played a role in communication between free-swimming single cells or between cells within a colonial life history phase. During the evolution of metazoans this pre-existing signalling machinery has been moved to cell-cell contact sites forming a classical synapse. In this scenario, the function and the biological role (cell-cell signalling) is ancestral and conserved feature but the localization in a synapse-like setting would have been an evolutionary novelty (Wagner 2018).
2. The components of neurosecretory vesicles have already been present in the last common ancestor of holozoans but were not involved in cell-cell signalling. Instead, they might have played a role in the secretion of substances such as ECM components or enzymes. In the last common ancestor of the Metazoa, these components have then been co-opted (Gould and Vrba 1982) for cell-cell signalling and moved to cell-cell contact sites. In this scenario, the function would be ancestral, while the biological role and localization would represent evolutionary novelties.

Our 3D reconstructions of two unicellular choanoflagellate cells revealed a polarized and diverse vesicular landscape. We acknowledge that vesicles are highly dynamic organelles and thus, it can be problematic to assign single vesicles to one of the gross types since they exhibit intermediate features (diameter, electron density, localization) between two types. The large whiskers in Figure 3L is a visualization of this problem. However, since either the median values or electron densities (or both) of the assigned vesicle types are very different from each other our described vesicle types are highly likely to be real. In the choanoflagellates *S. rosetta* and *M. brevicollis*, we identified morphologically distinct vesicle populations at the basal and apical pole of the cell. Both vesicle populations are potentially of secretory nature, as we found that the vesicle-associated SNARE protein synaptobrevin is localized to the apical and basal part of the choanoflagellate *S. rosetta*. We can only speculate about the content and function of these vesicles. Vesicles localized to the basal pole of *S. rosetta* could potentially contain the C-type lectin Rosetteless (Levin *et al*. 2014), or other extracellular matrix material (Wetzel *et al*. 2018), as their basal secretion seems to be essential for multicellular rosette development (Wetzel *et al*. 2018). Vesicles at the apical pole, close to the feeding collar in choanoflagellates, might contain mucus and digestive enzymes for external digestion (Arendt *et al*. 2015). Alternatively, these vesicles could also contain sialic acid, aspartate or glutamate (Arendt 2020), which they might use for communication between cells, as our comparative analysis revealed putative sialin-like transporters (Miyaji *et al*. 2008) in the genomes of *M. brevicollis* and *S. rosetta*.

Despite the lack of data on vesicular cargo and the function of vesicular signalling molecules in unicellular holozoans we propose a possible, choanoflagellate-biased scenario for the structural evolution of neurosecretory cell-cell signalling (Figure 4). Most of the structural components of neurosecretory vesicles as well as a polarized vesicular transport were present in the last common ancestor of choanoflagellates and Metazoa. In colonial choanoflagellates plasma membrane contacts were present but not involved in chemical cell-cell signalling. In the stem lineage of metazoans, neurosecretory vesicles were recruited to plasma membrane contact sites (soma or filopodial contacts) and used for intracolonial communication. This resulted in the emergence of a “presynaptic” (signal donor) and “postsynaptic” (signal receptor) cell. This condition might have been further stabilized in the “epithelialized” last common ancestor of the metazoans. From this condition many different structural types of synapses, like the presently described neuroid-choanocyte relationship in a sponge (Musser *et al*. 2019), presynaptic triads and somatic synapses in ctenophores (Hernandez-Nicaise 1968) and “classical” synapses present in most other metazoans might have evolved. In this scenario, the structural co-option of ancestral neurosecretory vesicles and polarized vesicular transport at plasma membrane contact sites might be the key process leading the structural evolution of metazoan synapses. However, data on the presence, intra-cellular localization and function of classical “metazoan” neurotransmitters in unicellular holozoans are needed to elucidate the ancestral function of the neurosecretory vesicle machinery.

**Figure 4.**
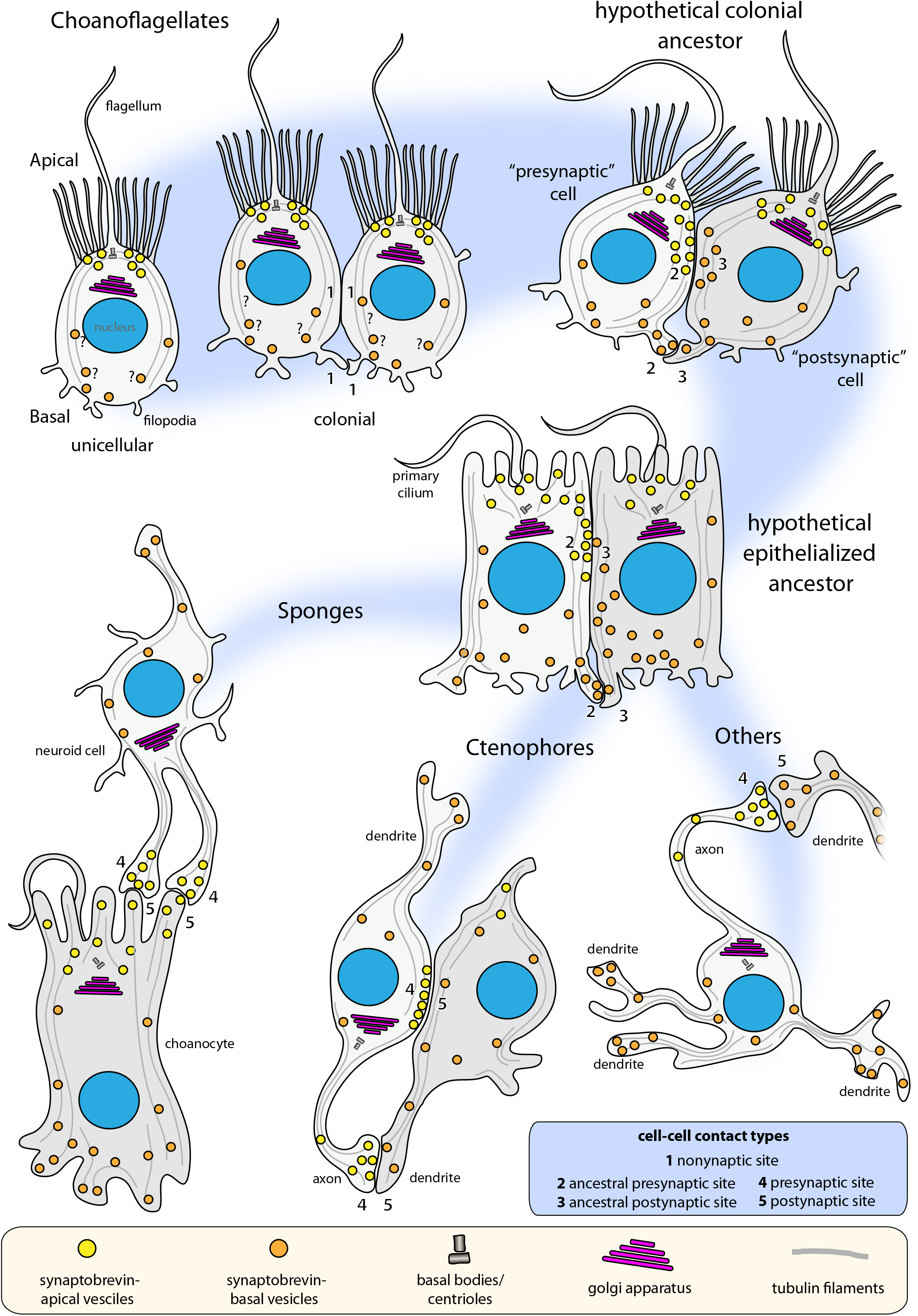
An evolutionary scenario for the structural evolution of metazoan synapses. A polarized vesicle transport system might have already existed in choanoflagellates. No chemical signal transduction appears at soma or filopodial plasma membrane contact sites (1). In the stem lineage of the Metazoa, the apical-basal directed vesicle transport has been translocated to soma and/or filopodial plasma membrane contact sites. This resulted in one cell acting as a signal donor or ancestral presynapse (2) and another cell acting as a signal receiver or ancestral postsynapse (3). This relationship might have been further stabilized in a more epithelialized, obligate multicellular metazoan ancestor. From this condition the evolution of more stable presynaptic (4) and postsynaptic (5) relationship was achieved in different groups of early branching metazoans

## Material & Methods

### Protein Searches and Analysis

Putative protein homologs of neurosecretory vesicle proteins were found using the basic local alignment sequence similarity search tool (BLASTp) (Altschul *et al*. 1997) at the National Center for Biotechnology Information (NCBI), National Human Genome Research Institute (NIH) and EnsemblMetazoa database. As queries we used protein sequences from *Homo sapiens* and *Rattus norvegicus* for the detection of neurosecretory vesicle proteins in seven other metazoans (*Danio rerio, Strongylocentrotus purpuratus, Drosophila melanogaster, Nematostella vectensis, Trichoplax adhaerens, Mnemiopsis leidyi* and *Amphimedon queenslandica*), three protists (two choanoflagellates (*Monosiga brevicollis* and *Salpingoeca rosetta*) and one filasterean (*Capsaspora owczarzaki*) and three fungi (*Batrachochytrium dendrobatidis, Rhizopus oryzae, Saccharomyces cerevisiae*). We performed protein homology-based searches with the default BLAST parameters and an e-value threshold of ≤1e^-6^. Initially obtained sequences were reciprocally searched against the NCBI protein database using BLASTp to verify results. Protein sequences of putative homologs were further analysed using the protein domain prediction programs Pfam (Punta *et al*. 2012) and SMART (Schultz *et al*. 1998, Letunic and Bork 2017). Protein accession numbers, domain composition and time points of the investigations are shown in the supplementary table 1.

### *Salpingoeca rosetta* and *Monosiga brevicollis* cell culture

Colony-free *S. rosetta* cultures (50818, American Type Culture Collection) were grown with co-isolated prey bacteria in 0.22 μm filtered choanoflagellate growth medium diluted at a ratio of 1:4 with autoclaved seawater as previously described (Laundon *et al*. 2019). Cultures were maintained at 18°C and split 1.5:10 once a week.

*M. brevicollis* cultures (50154; American Type Culture Collection) were cultured in artificial sea water mixed with Wards cereal grass medium in a 1:1 ratio, adjusted to a salt concentration of 53 mS/cm and sterile filtered as previously described (Burkhardt *et al*. 2011). Cultures were maintained at 25 °C and diluted 1:100 once a week.

### Protein Expression and Purification of *S. rosetta* synaptobrevin

To express and purify *S. rosetta* synaptobrevin a codon-optimised nucleotide sequence encoding the soluble portion of the protein [Syb (1-75)] was prepared by gene synthesis (Genscript, USA): GAGGCGAACCGTACCGGTGACTACCGTCTGCAGGAAGCGCAGCGTCAAGTGGGCGAAGTTCAAAACGTGATGCGTGATAACCTGACCAAGGTTATCGAGCGTGGTGAAAAACTGGACGATCTGGACGCGAAGGCGGAAGATCTGGAGGCGGAGGGTCAGCGTTTCCAAAACCGTGCGGGCCGTCTGCGTCGTCAGATGTGGTGGCAAAACAAACGTAACCAGTAA

This sequence was cloned into a pET28a(+) vector (69864, Novagen), which contains an N-terminal, thrombin-cleavable His6-tag. *Escherichia coli* BL21(DE3) Singles™ Competent Cells (70235, Novagen) were subsequently transformed. Following this, [Syb (1-75)] was expressed at 37°C for 3 h and purified by Ni^2+^-nitrilotriacetic acid (NTA) chromatography. For this, *E. coli* cells were pelleted at 3,488x g for 10 min at 4°C (Heraeus™ Megafuge™ 40R) and incubated at room temperature for 20 min with 100 μl of 200 mM phenylmethanesulfonyl fluoride (PMSF) (36978, ThermoFisher Scientific) and lysozyme from chicken egg white (L6876, Sigma-Aldrich). Following incubation, cells were sonicated (Vibra-CellTM, Sonics®) by 3 x 30 s pulses and incubated for 10 min at room temperature with 100 μl of 1 M MgCl2, 500 μl of 20% Triton X-100 and Deoxyribonuclease 1 from bovine pancreas (DN25, Sigma-Aldrich). Cellular debris was removed by centrifugation at 5,488x g for 40 min at 4°C (Heraeus™ Megafuge™ 40R) and incubated for 2 h with 500 μl HisPurTM Cobalt Resin (89964, ThermoFisher Scientific) at 4°C. The beads were then pelleted at 1,363x g for 10 min at 4°C, the supernatant was removed, and then washed three times in wash buffer (500 mM NaCl, 20mM Tris pH 7.4). His-tagged proteins were eluted from the beads in a disposable polypropylene column (29924, ThermoFisher Scientific) using elution buffer (4 ml wash buffer containing 400 mM imidazole) and dialysed overnight in Biodesign™ Cellulose Dialysis 3.5 kDa tubing (12757496, ThermoFisher Scientific) at 4°C in dialysis buffer (100 mM NaCl, 20mM Tris pH 7.4, containing 15 μl bovine thrombin (605157, Merck Millipore) to cleave the His-tags). Protein eluates were further purified by ion exchange chromatography using an Äkta Prime Plus equipped with a HiTrapTM SP HP column (GE Healthcare, Sweden) and eluted along a linear gradient of NaCl in 20 mM Tris, pH 7.4, 1 mM EDTA. The success of the purification was assessed by sodium dodecyl sulfate polyacrylamide gel electrophoresis (SDS-PAGE) on a 16% gel, run using an XCell SureLock™ MiniCell chamber (ThermoFisher Scientific, USA) and stained with SimplyBlue™ SafeStain (LC6060, ThermoFisher Scientific). Protein concentrations were quantified by measuring absorbance at 280 nm using a NanoDrop 1000 Spectrophotometer (ThermoFisher Scientific, USA).

### *Salpingoeca rosetta* synaptobrevin antibody production

Polyclonal antibodies were commercially raised in rabbits against recombinant [Syb (1-75)] antigen (Covalab, UK). Immunoglobulins were purified against Protein A from 1 ml of rabbit serum using a Protein A HP SpinTrap (28-9031-32, GE Healthcare) following the manufacturer’s instructions.

### Immunofluorescence Microscopy

Prior to fixation, cells were pelleted by gentle centrifugation (500x g for 10 min at 4°C) in a Heraeus™ Megafuge™ 40R (ThermoFisher Scientific) and resuspended in a small volume of culture medium. Concentrated cell suspension (500 μl) was applied to glass-bottom dishes coated with poly-L-lysine solution (P8920, Sigma-Aldrich) and left for 10-30 min until cells were sufficiently adhered. Cells were fixed in 200 μl of ice-cold 6% acetone in 4 x PBS for 5 min and then 4% paraformaldehyde in 4 x PBS for 15 min. Fixing solutions were then aspirated off, washed twice in 4 x PBS, twice in 2 x PBS and once in 1 x PBS and then blocked for 30 min in blocking buffer (1% bovine serum albumin (BSA) and 0.6% Triton X-100 in PEM solution (100 mM piperazine-N,N’-bis(2-ethanesulfonic acid) (PIPES) at pH 6.9, 1mM EGTA, and 0.1 mM MgSO4)). Cells were then incubated with primary antibodies (mouse monoclonal antibody against β-tubulin (E7, 1:200; Developmental Studies Hybridoma Bank, USA) and anti-[Syb(1-75)] antibody, 1:500) in 200 μl of blocking buffer for 1 h, washed 4 times in blocking buffer, and then incubated in the dark for 1 h with secondary antibodies in 200 μl of blocking buffer (polyclonal goat anti-mouse Alexa Fluor^®^ 488, 1:200 (A32723, ThermoFisher) and goat anti-rabbit Alexa Fluor^®^ 647, 1:200 (A-21244, ThermoFisher)). Dishes were then washed four times in blocking buffer, washed once in 1 x PBS and finally mounted under coverslips with ProLong^®^ Gold Antifade Mountant (P36935, ThermoFisher Scientific). Single choanoflagellate cells were imaged using a Zeiss LSM 510 confocal microscope. Colonial choanoflagellate cell were imaged using a Zeiss Axio Observer LSM 880 with an Airyscan detector.

### Transmission Electron Microscopy (TEM)

TEM of *S. rosetta* cells was performed essentially as described (Laundon *et al*. 2019). In brief, *S. rosetta* cells were high pressure frozen in a Bal-Tec HPM 010 high pressure freezer (Bal-Tec AG, Liechtenstein). Freeze substitution with 1% osmium tetroxide plus 0.1% uranyl acetate in acetone was performed over 2 h by the SQFS method of McDonald (McDonald and Webb 2011), then infiltrated with Eponate 12 resin and polymerized in a Pelco Biowave research microwave oven (Ted Pella, Inc., Redding, CA) over a period of 2 h. Sections were cut at 70 nm thickness, poststained with uranyl acetate and lead citrate, and viewed in a Tecnai 12 transmission EM (FEI Inc., Hillsboro, OR).

TEM of *M. brevicollis* cells was performed essentially as described (Burkhardt *et al*. 2011). In brief, for *M. brevicollis* electron microscopy, cells were flash-frozen in a Baltec HPM 010 high-pressure freezer. Cryosubstitution and embedding were performed in a Leica EM AFS. Cells were sequentially incubated at low temperature (−90 °C) in 0.1% tannic acid (100 h) and 2% OsO_4_ (7 h) in acetone. They were progressively brought to room temperature before being embedded in Epon (Electron Microscopy Sciences) and polymerized 24 h at 60 °C. Ultrathin 70 nm sections were cut and contrasted with uranyl acetate and lead citrate before being observed in a LEO 912 AB (Zeiss).

### 3D reconstruction and analysis

To better recognize thin membranous outlines of vesicles contrast was enhanced using the CLAHE plugin in Fiji (Zuiderveld 1994, Schindelin *et al*. 2012) for the image stack of *S. rosetta* prior to the reconstruction. Digital image stacks of the TEM sections of *M. brevicollis* and *S. rosetta* were imported into AMIRA (FEI Visualization Sciences Group) and aligned semi-manually. Subsequently, single vesicles were segmented manually by tracing structures along the z-axis and 3D reconstructed by automatically merging the traced parts. In some cases, there were fluent transitions between large vesicles and isolated smaller parts of the smooth endoplasmic reticulum. For consistent results membranous structures were defined as vesicles when they could be traced over three sections maximum and as part of the smooth endoplasmic reticulum when they were larger than three sections. For surface reconstructions single surface models for each vesicle were rendered from the segmented materials, numbers of vertices were reduced around ten times and the surfaces were smoothened. The cell soma, collar and flagellum were visualized and merged with the vesicle surface models using the volume rendering function in AMIRA. Vesicle diameters were calculated using the 3D measuring tool in AMIRA. For every vesicle, the largest distance between two points on the vesicular membrane, evaluated by eye, was measured. If vesicles extended over several (maximum three) sections, the diameter was measured on the section with the largest surface area. All measurements were conducted using unprocessed, unsmoothed materials. Subsequently, all measurements were exported to Microsoft Excel 2010 (Microsoft Corporation) to prepare point graphs and box and whisker plots.

## Acknowledgments

We thank Tarja Hoffmeyer and Maria Sachkova for valuable comments on the paper. This work was supported the Sars Centre core budget.

## Supplementary material

Supplementary table 1: Protein accession numbers and domain composition of neurosecretory proteins

Supplementary table 2: Vesicle diameters measured in a cell of *Salpingoeca rosetta* and *Monosiga brevicollis*.

Supplementary video 1: 3D vesicular landscape of the choanoflagellate *Salpingoeca rosetta*. Supplementary video 2: 3D vesicular landscape of the choanoflagellate *Monosiga brevicollis*.

## Additional Information

### Data Accessibility

The datasets supporting this article have been uploaded as part of the Supplementary Material.

### Authors’ Contributions

RG, BN and PB designed the story. RG performed the comparative protein analysis. BN reconstructed the choanoflagellates *S. rosetta* and *M. brevicollis*. DL expressed and purified synaptobrevin and performed immunostainings of colonial *S. rosetta*. PB performed immunostainings of single *S. rosetta*. CI, BC, FV performed TEM of *M. brevicollis*. KM performed TEM of *S. rosetta*. RG, BN, CI, KM, BC, FV, DF and PB analyzed data. RG, BN and PB wrote the manuscript.

